# Network Analysis Reveals Protein Modules Associated with Childhood Respiratory Diseases

**DOI:** 10.1101/2024.06.14.599044

**Authors:** Nicole Prince, Sofina Begum, Kevin M Mendez, Lourdes G Ramirez, Yulu Chen, Qingwen Chen, Su H Chu, Priyadarshini Kachroo, Ofer Levy, Joann Diray-Arce, Paolo Palma, Augusto A Litonjua, Scott T Weiss, Rachel S Kelly, Jessica A Lasky-Su

**Author notes:** **Corresponding Author:** Jessica A. Lasky-Su, ScD., Channing Division of Network Medicine Department of Medicine, Brigham and Women’s Hospital, 181 Longwood Avenue, Boston, MA, 02115. These authors contributed equally (Rachel S Kelly, Jessica A Lasky-Su). **Funding:** This study was funded by NHLBI grants R01HL123915, R01HL141826, K01HL146980, and T32HL007427; and NIAID grant U19AI168643.

## Abstract

**Background:** The first year of life is a period of rapid immune development that can impact health trajectories and the risk of developing respiratory-related diseases, such as asthma, recurrent infections, and eczema. However, the biology underlying subsequent disease development remains unknown.

**Methods:** Using weighted gene correlation network analysis (WGCNA), we derived modules of highly correlated immune-related proteins in plasma samples from children at age 1 year (N=294) from the Vitamin D Antenatal Asthma Reduction Trial (VDAART). We applied regression analyses to assess relationships between protein modules and development of childhood respiratory diseases up to age 6 years. We then characterized genomic, environmental, and metabolomic factors associated with modules.

**Results:** WGCNA identified four protein modules at age 1 year associated with incidence of childhood asthma and/or recurrent wheeze (P_adj_ range: 0.02-0.03), respiratory infections (P_adj_ range: 6.3×10-9-2.9×10-6), and eczema (P_adj_=0.01) by age 6 years; three modules were associated with at least one environmental exposure (P_adj_ range: 2.8×10-10-0.03) and disrupted metabolomic pathway(s) (P_adj_ range: 2.8×10-6-0.04). No genome-wide SNPs were identified as significant genetic risk factors for any protein module. Relationships between protein modules with clinical, environmental, and ‘omic factors were temporally sensitive and could not be recapitulated in protein profiles at age 6 years.

**Conclusion:** These findings suggested protein profiles as early as age 1 year predicted development of respiratory-related diseases through age 6 and were associated with changes in pathways related to amino acid and energy metabolism. These may inform new strategies to identify vulnerable individuals based on immune protein profiling.

## INTRODUCTION

The first 1000 days of life are critical to immune development, and disruptions to normal physiological processes during this period can have long-lasting health consequences, including increased risk of respiratory and related diseases, such as asthma, recurrent wheeze, eczema, and frequent infections.^1^ These impose high public health burdens,^2^ and while genetic and environmental risk factors have been identified, these cannot fully explain inter-individual variability in risk. Protein mediators such as cytokines, chemokines, and growth factors regulate immune responses^3^ and reflect variability in responses between individuals.^2^ Many of these proteins have established clinical relevance as biomarkers or therapeutic targets in respiratory diseases,^4^ and they have traditionally been studied with respect to single protein features. However, investigating protein profiles – rather than individual components – is an emerging conceptual framework that has provided insights into disease mechanisms.^5^ Applying this framework to study early life immune development could provide similar insights to characterize underlying biology leading to disease.

Plasma protein profiles in early life reflect the influence of genetic and environmental cues,^6^ but the impact of protein profiles in disease risk is incompletely understood. Several studies have demonstrated the contribution of genetic risk and early life environment on respiratory diseases,^7^ but their impacts are incompletely understood. Disruptions to normal immune function produce observable alterations to downstream biochemical pathways, reflected in the metabolome.^8^ These metabolomic changes can provoke further dysregulation related to disease progression.^9^ Investigating relationships between early life protein profiles with environmental factors and downstream metabolomic consequences could provide new insights into the antecedents and pathophysiology of childhood respiratory-related diseases. Further, defining these relationships with respect to groups of highly-correlated protein features, rather than individual protein targets, could enhance understanding of the complex biological milieu present during early life immune development, as these could suggest involvement in common or complementary processes affecting immune development in early life.

In this study, we sought to define relationships between immune protein profiles and childhood respiratory-related diseases, including asthma, recurrent wheeze, respiratory infections, and eczema. Our hypothesis was that network analysis would identify clinically-relevant modules of highly-correlated proteins in early life; further, these protein modules would be related to genetic, environmental, and metabolomic factors providing further biological insights. We utilized protein profiling data from children at age 1 in the Vitamin D Antenatal Asthma Reduction Trial (VDAART)^10^, then investigated associations between modules and clinical outcomes. Modules were further characterized with respect to the genome, environmental exposures, and metabolomic profiles to enhance molecular insights across multiple layers of systems biology.^11^

## METHODS

### Vitamin D Antenatal Asthma Reduction Trial (VDAART)

The Vitamin D Antenatal Asthma Reduction Trial^10,12^ was a clinical trial from 2009-2015 that recruited pregnant women between 10-18 weeks’ gestation (GW) and randomized them to a daily vitamin D dose of 4400 IU or 400 IU as normal pregnancy care. A subset of 294 mother-child pairs from VDAART were utilized in this study, based on availability of plasma sample for proteomic profiling. Detailed information on VDAART individuals can be found in the **Supplementary Methods** and in **Table 1**. Pregnant mothers completed monthly questionnaires throughout the duration of pregnancy, including questions about diet quality,^13^ smoking, and demographic/social characteristics. At delivery, birth characteristics were collected, including birth weight, gestational age at delivery, mode of delivery. Offspring of VDAART mothers were monitored over the first six years of life at yearly clinical visits and through quarterly questionnaires completed by parents/caregivers. At yearly visits, blood samples were collected from offspring. This study was approved by the Partners Human Research Committee at Brigham and Women’s Hospital (Protocol 2014P001109).

**Table 1.** Characteristics of VDAART Children at Age 1 Year. A subset of 294 children from the VDAART study with proteomic profiling were included in this study. Clinical outcomes, demographic characteristics, and other social and environmental exposure variables are reported below compared to all VDAART individuals with available clinical data.

|  | Individuals with Protein Data (N=294) | VDAART Overall (N=880) |
| --- | --- | --- |
| **Childhood Clinical Outcomes** |  |  |
| Reported Infections ages 0-6 years, Mean (SD) | 28.45 (12.94) | 28.54 (13.30) |
| Asthma and/or Wheeze Diagnosis ages 0-6 years, N (%) | 135 (45.9) | 359 (44.6) |
| Recurrent Wheeze Diagnosis ages 0-6 years, N (%) | 129 (43.9) | 343 (42.6) |
| Eczema Diagnosis ages 0-6 years, N (%) | 151 (52.8) | 369 (52.3) |
| **Demographic Characteristics** |  |  |
| Child sex, N male (%) | 151 (54.1) | 254 (54.2) |
| Reported Race Category, N (%) |  |  |
| White | 89 (31.9) | 152 (32.4) |
| Black | 134 (48.0) | 227 (48.4) |
| Other | 56 (20.1) | 90 (19.2) |
| Study Site |  |  |
| Boston, MA, N (%) | 89 (31.9) | 144 (30.7) |
| San Diego, CA, N (%) | 93 (33.3) | 158 (33.7) |
| St. Louis, MO, N (%) | 97 (34.8) | 167 (35.6) |
| Reported Annual Household Income, N (%) |  |  |
| Less than \$30,000 | 82 (39.8) | 263 (39.3) |
| \$30,000-\$49,999 | 37 (18.0) | 120 (17.9) |
| \$50,000-\$74,999 | 32 (15.5) | 101 (15.1) |
| \$75,000-\$99,999 | 25 (12.1) | 82 (12.3) |
| \$100,000-\$149,999 | 22 (10.7) | 71 (10.6) |
| Over \$150,000 | 8 ( 3.9) | 32 ( 4.8) |
| Reported Maternal Education Level, N (%) |  |  |
| Did Not Graduate High School | 36 (12.9) | 108 (12.3) |
| High School Graduate | 67 (24.0) | 225 (25.6) |
| Technical School, Junior College, or Some College | 82 (29.4) | 256 (29.1) |
| College Graduate | 57 (20.4) | 165 (18.8) |
| Graduate School or Higher | 37 (13.3) | 126 (14.3) |
| **Prenatal Exposures** |  |  |
| Maternal Vitamin D at 10-18GW in ng/mL, Mean (SD) | 23.03 (9.76) | 22.80 (10.25) |
| Maternal Vitamin D at 32-38GW in ng/mL, Mean (SD) | 33.38 (14.47) | 32.90 (14.63) |
| Exposure to smoking during pregnancy, N (%) | 76 (27.3) | 234 (31.1) |
| **Perinatal Exposures** |  |  |
| Cord Blood Vitamin D in ng/mL, Mean (SD) | 24.31 (11.82) | 23.16 (11.73) |
| Birth weight in kg, Mean (SD) | 3.31 (0.50) | 3.30 (0.53) |
| Gestational age at delivery in weeks, Mean (SD) | 39.20 (1.47) | 38.91 (2.27) |
| C-section mode of delivery, N (%) | 78 (28.0) | 241 (29.6) |
| Birth order, Mean (SD)<sup>1</sup> | 0.92 (1.04) | 0.91 (1.06) |
| **Postnatal Exposures** |  |  |
| Child Vitamin D at age 1 year in ng/mL, Mean (SD) | 29.86 (10.34) | 29.65 (10.32) |
| Child Body Mass Index (BMI) at age 1 year, Mean (SD) | 17.45 (2.26) | 17.43 (2.15) |
| Attended daycare between ages 0-1 year, N (%) | 31 (11.1) | 46 ( 9.8) |
| Breastfeeding Duration, N (%) |  |  |
| No breastfeeding | 51 (18.3) | 106 (22.6) |
| Breastfed < 6 months | 115 (41.2) | 184 (39.2) |
| Breastfed 6-12 months | 43 (15.4) | 80 (17.1) |
| Breastfed > 12 months | 70 (25.1) | 106 (22.6) |

<sup>1</sup>Birth order is derived from the number of children previously birthed by the mother

### Clinical Outcomes

A detailed description of asthma, recurrent wheeze, eczema, and infection definitions have been reported previously for VDAART, and these outcomes were collected through quarterly questionnaires.^12,14^ Caregiver report of: (i) doctor’s diagnosis of asthma and/or recurrent wheezing any time between birth and age 6 years, (ii) caregiver report of the cumulative number of respiratory infections during the first 6 years of life, and (iii) caregiver report of doctor’s diagnosis of eczema any time between birth and age 6 years.

### Proteomic, Genomic, and Metabolomic Profiling

Non-fasting blood samples were collected from mothers during pregnancy visits (10-18 GW; 32-38 GW), from cord blood at delivery, and from offspring at follow-up visits. Maternal blood samples were assayed for 25OHD levels. In plasma samples collected in offspring at ages 1 and 6 years, 200 immune-mediating proteins were measured using an available targeted NULISA-Seq panel that employs sequential immunocomplex capture and release, then next generation sequencing (NGS) to provide ultra-high sensitivity multiplexing (Alamar Biosciences, Freemont, CA, USA).^15^ A full list of protein targets can be found in **Supplementary Table 1**. Samples were selected for protein profiling based on existing metabolomic profiling data and availability of remaining plasma sample. Genotyping was performed in children using the Illumina Infinium HumanOmniExpressExome Bead chip (San Diego, CA, USA), as described previously.^16^ Metabolomic profiling of offspring plasma samples was performed by Metabolon, Inc. (NC, USA) using Metabolon’s global platform that generates data using High Performance Liquid Chromatography coupled to tandem Mass Spectrometry (HPLC-MS/MS).^14,17^ Additional details of proteomic, genomic, and metabolomic profiling can be found in the Supplementary Methods.

### Protein Module Generation Using Weighted Gene Correlation Network Analysis (WGCNA)

WGCNA^18^ was used to derive modules of highly-correlated proteins at age 1 year based on pairwise correlations between protein features using the WGCNA package in R v4.3.0.^19^ Modules were merged using a cut height (i.e., the Euclidean distance between modules) of 0.3 and a soft power threshold of 6 based on iterative process to identify an optimal number of modules. Following WGCNA, protein groups (i.e., all protein features within a respective module) were input into the STRING database version 12.0^20^ to identify common biological functions, and module hubs were identified based on the greatest number of edges. Modules were summarized as eigenvectors based on the first principal component of the included proteins for each individual. This eigenvector value for each module was utilized in subsequent statistical models to estimate associations between modules and clinical outcomes, ‘omics, and social/environmental characteristics. Age 1 modules were recapitulated using protein profiling from age 6 samples to assess the consistency of the module relationships with our outcomes if interest over time.

### Statistical Analysis

Associations with asthma, recurrent wheeze, and eczema outcomes were evaluated using logistic regression, in which outcomes were defined as “true” if any incidence was reported by caregivers at any point between birth and age 6 years. Poisson regression was utilized for the count of cumulative respiratory infections, which was represented by a continuous variable. In both cases, protein module eigenvalues were treated as predictors. Fully adjusted models included sex (1=male; 0=female), race (“White” used as reference group), breastfeeding duration (0=>12 months; 1=6-12 months; 2=<6 months; 3=none reported), and daycare attendance between birth and age 1 (true/false for any attendance).

Associations between protein module eigenvalues and social, environmental, and demographic variables were also evaluated, specifying module eigenvalues as outcomes. Logistic regression models estimated associations between modules and sex, smoking in pregnancy, mode of delivery, and daycare attendance. Linear models were used for reported annual household income, maternal education, race, breastfeeding duration, maternal diet quality scores, birth weight, birth order, age 1 BMI, and vitamin D levels in nanograms per milliliter (ng/mL). Correlations between these variables were estimated using the cor package in R v4.3.0.

Genome-wide associations between protein modules and single nucleotide polymorphisms (SNPs) were investigated using an expression quantitative trait loci (eQTL) approach using the Matrix eQTL package in R.^21^ Linear regression models estimated associations between protein module eigenvalues and individual metabolites at age 1 year using the glm package in R v4.3.0. Beta estimates and P-values from these models were used as input for metabolomic set enrichment analysis based on KEGG annotations using MetaboAnalyst v6.0.^22^

All regression analyses employed multiple testing correction controlling for false discovery rate (FDR) using the Benjamini-Hochberg procedure.^23^ An overview of the study design and statistical analysis procedures can be found in Figure 1.

**Figure 1.**
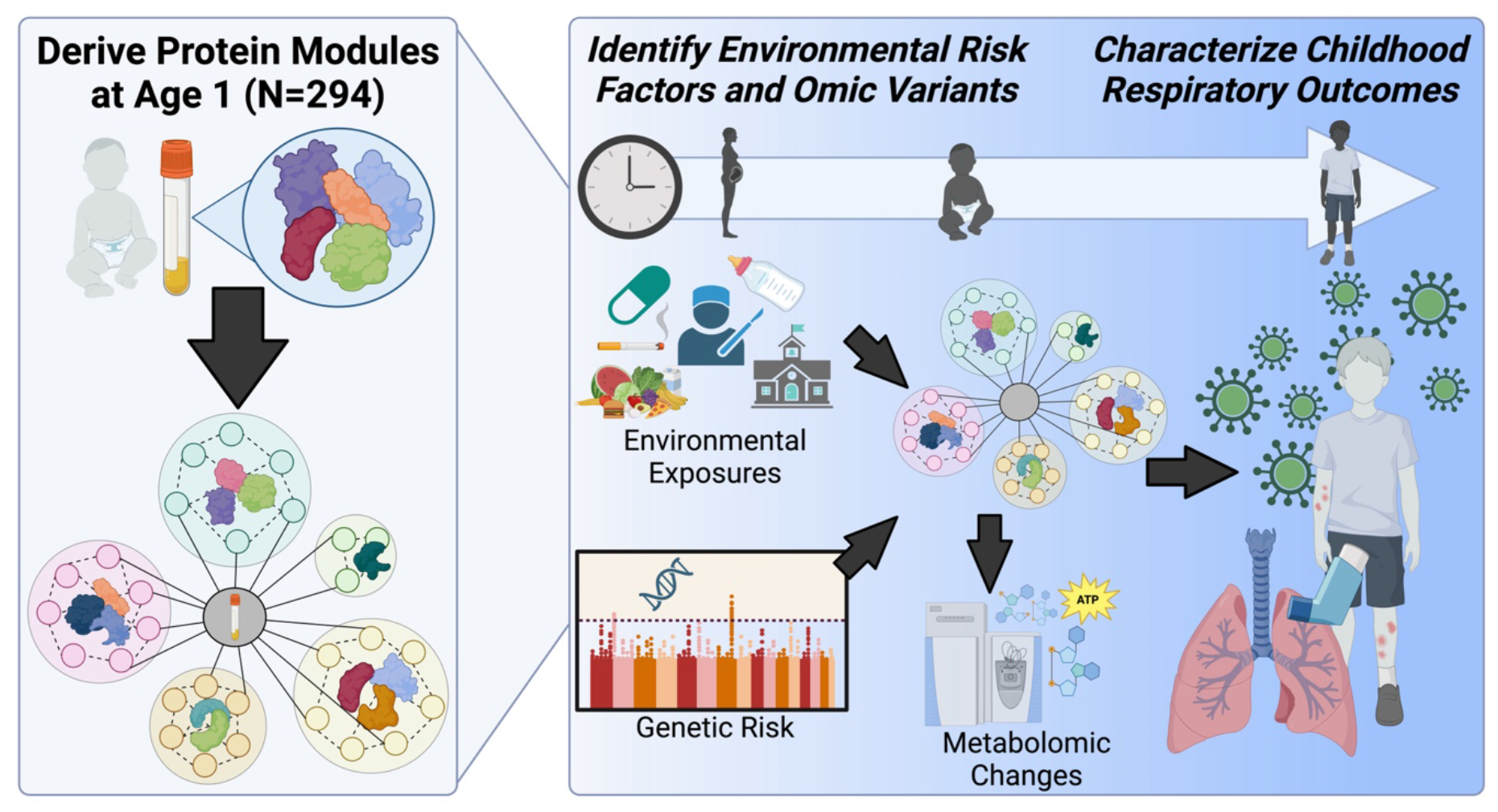
Overview of Approach. Plasma samples were collected from 294 children aged 1 and 6 years in the Vitamin D Antenatal Asthma Reduction Trial (VDAART), and targeted protein profiling of 200 immune-mediating proteins was performed using the NULISA-Seq platform from Alamar Biosciences, Inc. Using Weighted Gene Correlation Network Analysis (WGCNA), we derived protein modules based on correlations between plasma protein levels at age 1. After determining protein modules, we characterized module associations with respiratory-related outcomes, including incidence of asthma, recurrent wheeze, infections, and eczema by age 6 years. Genetic, environmental, and metabolomic factors related to outcomes were assessed to provide additional biological insights.

## RESULTS

### VDAART Offspring Clinical Outcomes

Among individuals selected for proteomic profiling, 45.9% had asthma, 43.9% had recurrent wheeze, and 52.8% had eczema by age 6. On average, children experienced 28.5 infections in the first 6 years, as reported by caregivers (Table 1). The subset of VDAART children included in this study was heterogeneous in demographic characteristics such as race (48.0% Black, 31.9% White and 20.1% Other), geographical location (31.9% in Boston, MA; 33.3% in San Diego, CA; 34.8% in St. Louis, MO), and maternal education status (33.7% reported bachelor’s degree or higher). A substantial proportion of VDAART participants reported low household income levels, with 39.8% of families reporting less than $30,000 per year. The subset of 294 individuals with protein profiling included in this study was representative of the overall VDAART cohort with respect to outcomes, demographic characteristics, and environmental exposures of interest (Table 1).

### Protein module associations with clinical outcomes were time-sensitive

WGCNA produced seven total protein modules based on protein profiles at age 1 year, four of which were associated with clinical outcomes by age 6 years using an FDR-adjusted P-value cutoff of 0.05 (**Fig. 2a-b**). All P-values reported in results are P-values after FDR correction, unless otherwise specified. Higher eigenvalues for Module 1 and Module 2 at age 1 were associated with higher incidence of asthma (Beta[CI]_Module1_= 5.8[1.6, 9.9], P_Module1_=0.03; Beta[CI]_Module2_= 5.2[1.1, 9.2], P_Module2_=0.03), recurrent wheeze (Beta[CI]_Module1_= 5.7[2.5, 9.8], P_Module1_=0.03; Beta[CI]_Module2_=4.7[0.7, 8.7], P_Module2_=0.03), and cumulative infections (Beta[CI]_Module1_= 1.2[0.8, 1.7], P_Module1_=6.3×10^-9^; Beta[CI]_Module2_= 1.0[0.6, 1.4], P_Module2_=2.9×10^-6^), while Module 3 was the only module associated with higher incidence of eczema (Beta[CI]_Module3_= 6.4[2.2, 10.6], P_Module3_=0.01). Module 4 demonstrated the opposite directions of effect with higher eigenvalues associated with reduced incidence of asthma (Beta[CI]_Module4_= -4.8[-8.9, -0.7], P_Module4_=0.03) and recurrent wheeze (Beta[CI]_Module4_= -4.9[-9.0, -0.8], P_Module4_=0.03) (**Fig. 2a**). Higher eigenvalues for all modules were correlated with elevated levels of individual proteins within each respective module (**Supplementary Table S1**). These results were unchanged after adjusting for sex, race, breastfeeding duration, and daycare attendance (**Supplementary Table S2**). These protein modules could not be recapitulated in protein profiles at 6 years (**Fig. 2b**); only the association between Module 2 at 6 years and cumulative infections retained significance (Beta[CI]_Module2_= 0.6[0.3, 1.01], P_Module2_=1.1×10^-3^). Protein correlations with module groupings at 6 years can be found in **Supplementary Table S3**.

**Figure 2.**
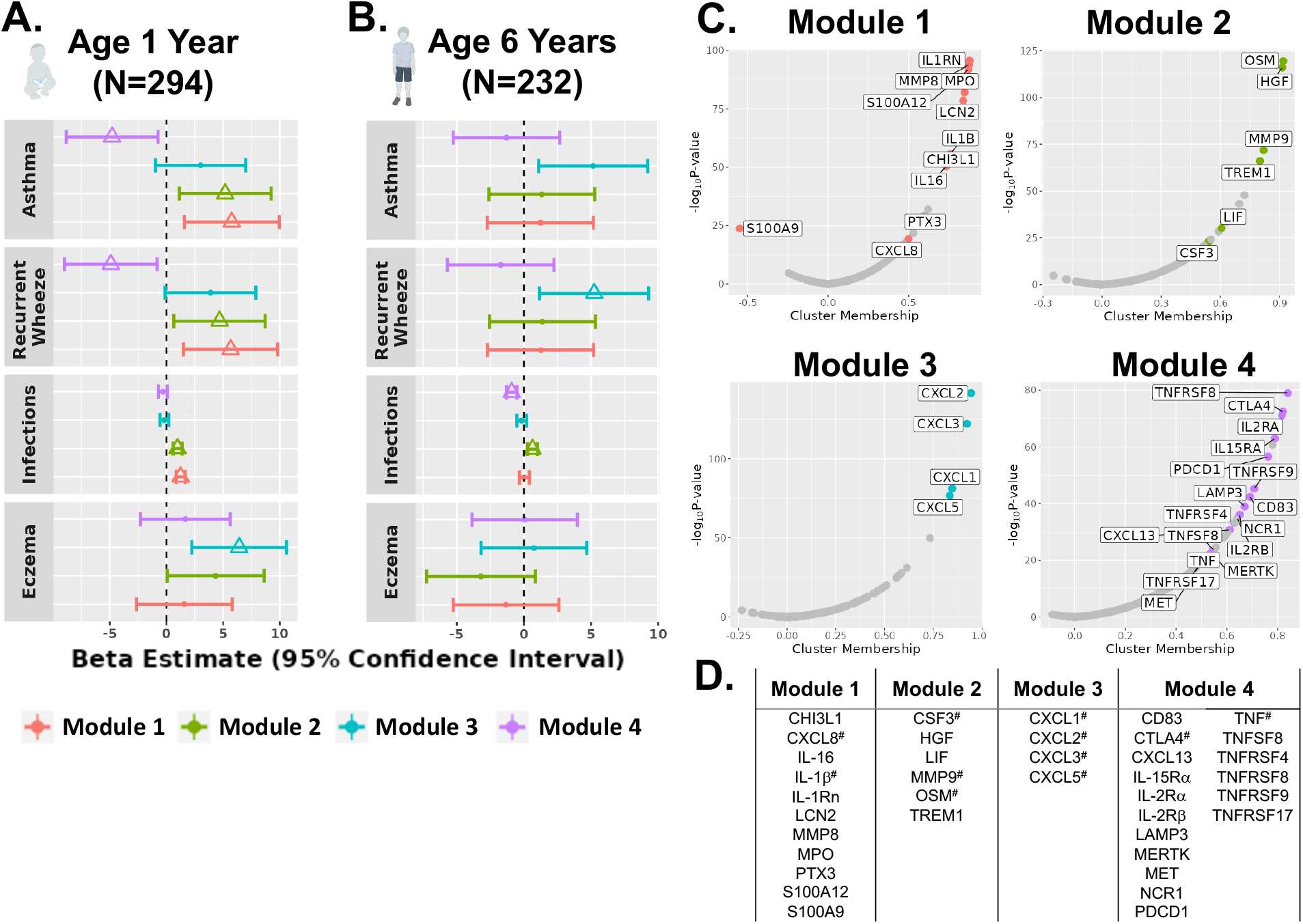
WGCNA identified four time-sensitive protein modules associated with clinically-relevant outcomes. Four modules were associated with outcomes in a temporally-sensitive manner; associations between modules and clinical outcomes are shown at age 1 year (A) and age 6 years (B). Beta and 95% confidence interval are displayed for each forest plot and colored by module; triangle point shape indicates the association was significant at an adjusted P-value < 0.05. (C) Correlations between individual proteins and Module 1, Module 2, Module 3, and Module 4 plotted in volcano plots. (D) Shows a list of individual protein features sorted into each module by WGCNA in alphabetical order, and module hubs are denoted with a hash mark (#).

Shared biological functions of proteins within modules were characterized using the STRING database (**Supplementary Figure S1**), and correlations between individual protein levels at age 1 and module eigenvalues were assessed. Module 1 included eleven proteins (**Fig. 2c**); interleukin-1 receptor (IL-1Rn) demonstrated the largest correlation (r^2^=0.88) with Module 1, followed by matrix metallopeptidase 8 (MMP8; r^2^=0.88). However, IL-1β and C-X-C Motif Chemokine Ligand 8 (CXCL8; also known as IL-8) showed the largest number of edges in the STRING network (i.e., connections between individual proteins based on publicly available sources of protein–protein interaction information) and were considered module “hubs” (**Supplementary Figure S1a**). Module 2 included six proteins (**Fig. 2c**), and its eigenvalue showed the strongest correlations with oncostatin M (OSM; r^2^=0.92) and hepatocyte growth factor (HGF; r^2^=0.91); OSM, glycoprotein colony-stimulating factor 3 (CSF3), and MMP9 showed the largest number of edges and were module hubs (**Supplementary Figure S1b**). Module 3 included only four chemokines with all correlations above 0.8 (**Fig. 2c**): chemokine ligand 1 (CXCL1; r^2^=0.85), CXCL2 (r^2^=0.94), CXCL3 (r^2^=0.92), and CXCL5 (r^2^=0.83) with the module eigenvalue; all four proteins demonstrated an equal number of edges (**Supplementary Figure S1c**). Module 4 included the largest number of individual proteins (**Fig. 2c**), with TNF Receptor Superfamily Member 8 (TNFRSF8, r^2^=0.84), cytotoxic T-lymphocyte associated protein 4 (CTLA4, r^2^=0.82), and IL-2Ra (r^2^=0.82) demonstrating the highest correlations with module eigenvalue; CTLA4 and TNF were module hubs (**Supplementary Figure S1d**).

### Protein module associations with environmental variables and ‘omics

Environmental variables related to social determinants of health across the prenatal, postnatal, and demographic categories were associated with at least one protein module at a P-value threshold of 0.05 after multiple testing correction by FDR (**Fig. 3a**). However, the majority of associations with these variables was present only for Module 3, suggesting higher levels of Module 3 proteins in children were associated with maternal smoking (Beta[CI]= 9.16[4.3, 14.0]; P=9.1×10^-4^), non-White race (Beta[CI]= 5.7[4.1, 7.4]; P=2.8×10^-10^), lower maternal education (Beta[CI]= 4.1[1.7, 6.5]; P=4.4×10^-3^), and lower household income (Beta[CI]= 6.2[2.8, 9.7]; P=2.2×10^-3^). Module 3 was associated with poor maternal diet during the first (Beta[CI]= -6.3[-9.6, -2.9]; P=1.09×10^-3^) and third trimester (Beta[CI]= -7.2[-10.7, -3.6]; P=3.6×10^-4^) and lower vitamin D levels during the first trimester (Beta[CI]= -3.1[-5.1, -1.2]; P=6.9×10^-3^). VDAART was originally part of a vitamin D supplementation study, but only one significant association with vitamin D levels at any time period was observed (**Supplementary Table S4**).

**Figure 3.**
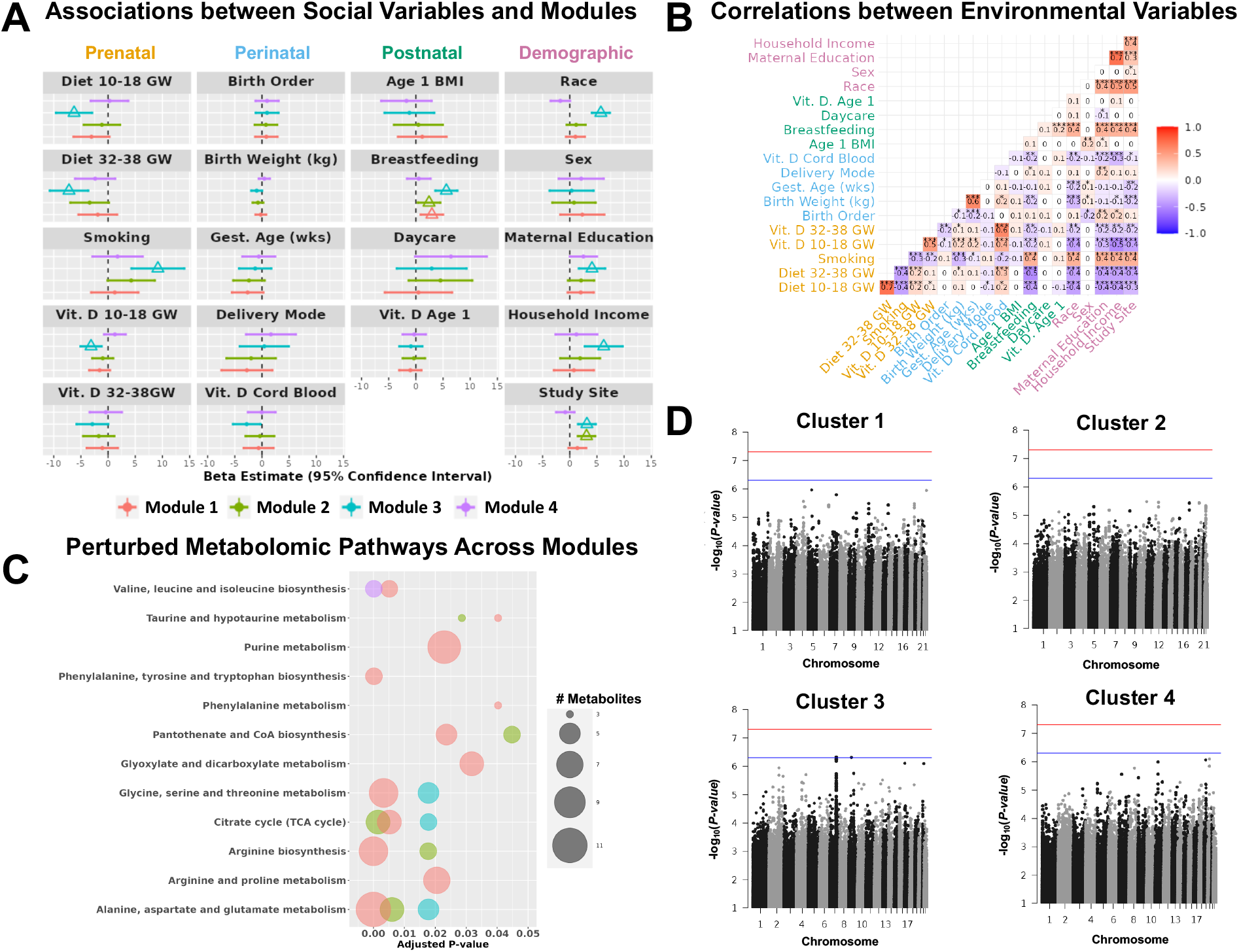
Protein modules were associated with other ‘omics. (A) Forest plots depict associations between environmental, social, and demographic variables and age 1 modules. Beta and 95% confidence intervals are shown colored by module. Associations meeting a significance threshold of adjusted P-value < 0.05 are denoted with large, open triangle shape. (B) Correlations between environmental, social, and demographic variables in all VDAART children at age 1 year (N=294); correlation coefficient is shown, and significance is noted by asterisks for P<0.05*, P<0.01**, and P<0.001***. Axis labels are colored by the relevant time period: prenatal (orange), perinatal (light blue), postnatal (forest green), and demographic (pink). (C) Bubble plot displaying enriched metabolomic pathways for each module at age 1 year; size of circle corresponds to the number of metabolites in each pathway, and coloring by module is consistent with panel (A). (D) Genome-wide associations with each age 1 protein module. Genome-wide significance threshold of 5×10^-8^ is marked by a red line and relaxed threshold of 5×10^-7^ is marked by a blue line.

Breastfeeding duration and study site were the only variables associated with multiple modules. Shorter breastfeeding duration was associated with higher levels of proteins in Module 1 (Beta[CI]= 2.9[0.9, 5.0], P=0.01), Module 2 (Beta[CI]= 2.4[0.3, 4.5], P=0.03), and Module 3 (Beta[CI]= 5.6[3.6, 7.6]; P=6.1×10^-7^). Study site was associated with Module 2 (Beta[CI]= 3.1[1.5, 4.9], P=3.9×10^-4^) and Module 3 (Beta[CI]= 3.1[1.5, 4.8]; P=3.9×10^-4^) in a direction that suggested location at urban centers located in Boston, MA and St. Louis, MO were associated with higher levels of proteins within these 2 modules compared to San Diego, CA. Correlations between environmental variables demonstrated a number of significant correlations (**Fig. 3b**) that should be considered when interpreting the influence of any individual factor.

Individual metabolites were associated with Module 1 (N=255 metabolites), Module 2 (N=150 metabolites, Module 3 (N=149 metabolites), and Module 4 (N=83 metabolites), and all associations between metabolites and modules are shown in **Supplementary Table S5**. The results of the *MetaboAnalyst* enrichment was based on KEGG pathways (**Fig. 3c**); Module 1 demonstrated the largest number of enriched pathways, with a total of 12 at an FDR-significant threshold (P-values=2.8×10^-6^ to 0.04) followed by Module 2 with 5 enriched pathways (P-values=1.49×10^-3^ to 0.04), then Module 3 with 3 enriched pathways (P-values=0.02), and finally Module 4 with only 1 enriched pathway (P-value=1.39×10^-4^). All metabolomic pathway enrichment results are available in **Table 2**. Genetic risk factors were also evaluated, but no genome-wide significant associations between SNPs and protein modules at age 1 year were observed based on a P-value threshold of 5×10^-8^; only two SNPs demonstrated associations with any module at a relaxed threshold of 5×10^-7^: rs6465878 of *FBXL13* and rs4878832 of *ANKRD18A* with Module 3 (**Fig. 3d-g**).

**Table 2.** Enriched Metabolomic Pathways for Protein Modules at 1 Year of Age. Regression estimates and P-values from associations between individual metabolites and protein modules at age 1 year were used as input for *MetaboAnalyst* software to perform metabolite set enrichment analysis using KEGG pathway assignments. The number of metabolites enriched in each pathway for each protein module is shown alongside the FDR-corrected P-value calculated by *MetaboAnalyst*. Pathways with an FDR<0.05 are shown in the table.

| Metabolomic Pathway | Protein Module | Number of Metabolites | Adjusted P-value |
| --- | --- | --- | --- |
| Valine, leucine and isoleucine biosynthesis | Module 1 | 4 | 0.00514 |
| Valine, leucine and isoleucine biosynthesis | Module 4 | 4 | 0.000139 |
| Taurine and hypotaurine metabolism | Module 1 | 3 | 0.0403 |
| Taurine and hypotaurine metabolism | Module 2 | 3 | 0.0286 |
| Purine metabolism | Module 1 | 10 | 0.0229 |
| Phenylalanine, tyrosine and tryptophan biosynthesis | Module 1 | 4 | 0.000172 |
| Phenylalanine metabolism | Module 1 | 3 | 0.0403 |
| Pantothenate and CoA biosynthesis | Module 1 | 5 | 0.0236 |
| Pantothenate and CoA biosynthesis | Module 2 | 4 | 0.0448 |
| Glyoxylate and dicarboxylate metabolism | Module 1 | 6 | 0.0318 |
| Glycine, serine and threonine metabolism | Module 1 | 8 | 0.0033 |
| Glycine, serine and threonine metabolism | Module 3 | 5 | 0.0178 |
| Citrate cycle (TCA cycle) | Module 1 | 6 | 0.00514 |
| Citrate cycle (TCA cycle) | Module 2 | 6 | 0.00149 |
| Citrate cycle (TCA cycle) | Module 3 | 4 | 0.0178 |
| Arginine biosynthesis | Module 1 | 8 | 3.18e-06 |
| Arginine biosynthesis | Module 2 | 4 | 0.0177 |
| Arginine and proline metabolism | Module 1 | 7 | 0.0205 |
| Alanine, aspartate and glutamate metabolism | Module 1 | 11 | 2.75e-06 |
| Alanine, aspartate and glutamate metabolism | Module 2 | 6 | 0.006 |
| Alanine, aspartate and glutamate metabolism | Module 3 | 5 | 0.0178 |

## DISCUSSION

In this study we integrated clinical and demographic data from a well annotated longitudinal cohort of infants with genotyping, targeted proteomics, and metabolomics to identify signatures correlating with clinical outcomes. Identifying early life contributors to respiratory disease development can be challenging, as immune development during this period results in complex biological environments. While many efforts have sought to better characterize factors leading to asthma,^16^ recurrent infections,^24^ and eczema^25^ during childhood, the biological overlap in these conditions is not fully understood. Network-based approaches have demonstrated utility in identifying important components of ‘omic data sets, and we applied this framework to profiles of immune-mediating proteins at age 1 year to identify protein modules most influential in each condition, as well as the overlap of multiple childhood respiratory-related diseases. Our investigation identified two protein modules associated with the overlap of asthma and recurrent infections and two additional protein modules specifically associated with asthma and with eczema, respectively. Module associations with outcomes were temporally sensitive, which highlights the importance of the first year of life in respiratory health trajectories. These findings may ultimately aid in pinpointing relevant biology related to early life immune development and its connection to respiratory diseases in childhood.

Our results demonstrated overlap between protein Modules 1 and 2 with childhood respiratory diseases, including asthma, wheeze, and infections. Module 1 captured proteins central in acute phase immune response;^26^ specifically, IL-1β and CXCL8- two potent pro-inflammatory factors-were Module 1 hubs,^27^ suggesting time-sensitive disruptions in acute phase response in early life may contribute to development of respiratory diseases. Proteins in Module 1 demonstrated the highest degree of metabolomic enrichment, including 12 pathways across amino acid and energy metabolism,^28^ which may reflect subsequent biochemical changes in response to the actions of proteins within this module. Proinflammatory and profibrotic mediators,^29,30^ and showed consistent relationships with outcomes, further implicating innate immune mechanisms in early life. Module 2 hubs CSF3, MMP9, and OSM participate in IL-6-type signaling that ultimately impacts a range of immune functions.^30^ Metabolomic enrichment of 5 pathways was consistent across Modules 1 and 2, which may suggest shared metabolomic changes associated with proteins within these modules. Reduced breastfeeding, a recognized risk factor in asthma^31^ and infections,^24^ was also associated with higher levels of proteins in Modules 1 and 2; no other genetic or environmental variables were identified, including vitamin D levels during pregnancy or in offspring.

In contrast, reduced levels of proteins in Module 4 at age 1 year were associated with development of asthma and recurrent wheeze. Module 4 proteins were broadly related to T cell regulatory processes, and the hubs of this module-CTLA4 and TNF-are regulators of T cell activation.^32^ Our findings suggested that reduced levels of these proteins, and thereby reduced regulation of T cell inflammation, may lead to increased risk of childhood asthma and wheeze. In addition to TNF as a Module 4 hub, we noted elevation of five TNF superfamily proteins known to participate in regulating the immune responses of T cells;^33^ with established connections to CXCL13^33^, further suggesting that lack of T cell regulation in early life may impact subsequent asthma risk. This was the only module with no associations to environmental or genetic variables. While asthma researchers have uncovered multiple genetic and environmental risk factors for disease,^16,34,35^ our results suggested that the resultant inflammatory profile represented by Module 4 proteins is not induced by a single genetic or environmental factor. Further, a single metabolomic pathway-valine, leucine and isoleucine biosynthesis-was enriched in this module, and the lowest number of individual metabolite associations were observed for Module 4.

Module 3 was the only module associated with childhood eczema and was comprised of four neutrophil attractant chemokines.^36^ CXCL1 and CXCL5 have been implicated in neutrophil chemotaxis to the skin during inflammatory episodes prevalent in atopic diseases,^37^ and our findings suggested increased neutrophilic inflammation is present in early life among infants at high risk of developing eczema, even outside of specific inflammatory episodes concomitant with disease. Enrichment of glycine, alanine, and TCA cycle pathways were also observed for Module 3, which could reflect the increased demand for energy sources in children with higher levels of inflammation mediated by these proteins.^38^ Importantly, prenatal and demographic variables were highly influential, so the temporal relationship between environmental characteristics and neutrophilic inflammation was confounded. Social determinants of health can greatly influence immune development in early life,^7^ and our study observed associations between elevated neutrophil chemokines of Module 3 and poor socioeconomic status. Additionally, lower vitamin D levels in early pregnancy were associated with higher levels of Module 3 proteins. While our study does not distinguish whether these are cause-effect relationships, these may represent relevant biomarkers to identify eczema-susceptible children at a very early time period.

One strength of this study is the availability of longitudinal clinical, demographic, environmental and systems biology data captured in VDAART,^12^ enabling identification of biomarkers and modules across multiple ‘omics that may influence clinical outcomes. Further, our protein profiling platform included broad coverage of immune-related proteins and was agnostic to specific immune mechanisms, allowing discovery across multiple immune-relevant pathways. However, relative quantification limited our ability to fully evaluate the time-sensitive impacts, especially given the dynamic nature of the immune system during early life.^39^ Fully quantitative profiling could enhance the clinical applicability of our findings and will be pursued in future studies. Additionally, the exploratory nature of this study necessitated a large amount of data, presenting a challenge in replication. Following the identification of particular protein groups, metabolomic pathways, and environmental exposures presented here, future studies will attempt replication in external populations to validate specific findings and explore relationships over a broader timespan.

In summary, this study demonstrated the utility of applying network analysis approaches to characterize biology in early life related to childhood respiratory-related diseases. Our results emphasized the importance of early life protein profiles in the development of asthma, eczema, and recurrent infections. The protein module hubs or modules themselves could represent novel biomarkers to identify children susceptible to these diseases, ultimately allowing earlier identification to combat these diseases in a vulnerable population.

## Supporting information

Supplementary Methods and Results

## ACKNOWLEDGEMENTS

We thank the participants and study staff of the original VDAART study and the team at Alamar Biosciences, Inc. for their assistance in quality control of the protein data acquired and used for this study.

