## Supplementary Methods and Results for "Network Analysis Reveals Protein Modules Associated with Childhood Respiratory Diseases"

### VDAART Cohort Details

#### *Collection of Demographic and Social Characteristics*

Demographic characteristics related to socioeconomic determinants were collected, including race, maternal education level, and household income. VDAART recruited across three clinical sites in the United States: Boston Medical Center, Boston; Washington University at St. Louis, St. Louis; and Kaiser Permanente Southern California Region, San Diego.[1] Mothers returned at 32-38 GW for a follow-up appointment. Offspring of VDAART mothers were monitored throughout the first 6 years of life via yearly clinical visits and quarterly questionnaires completed by parents/caregivers.[2]

#### *NULISA-Seq Protein Measurement*

Protein profiling data in plasma was performed by Alamar Biosciences, Inc (Fremont, CA, USA). Briefly, NULISA-Seq employs a multiplexed, sandwich immunoassay approach, as reported in detail previously.[3] A panel of 200 proteins was used in this study, including 124 cytokines, chemokines, and other proteins involved in inflammatory processes are included in this panel to capture a broad range of immune-related processes. This technology leverages immunocomplex capture of antigens to achieve attomolar sensitivity. Ligated reported oligonucleotides are formed between the capture and detection antibodies to form an immunocomplex specific to each protein target; this sequence is amplified by polymerase chain reaction (PCR); then, protein levels are reported using next generation sequencing (NGS) based on a library of pooled reporter molecules. In this study, the data produced relatively quantified data based on signal for each reporter molecular.

#### *Metabolomic Profiling*

Global metabolomic profiling was performed in plasma samples by Metabolon, Inc. (Research Triangle Park, NC, USA). All methods utilized a Waters ACQUITY ultra-performance liquid chromatography (UPLC) and a Thermo Scientific Q-Exactive high resolution/accurate mass spectrometer interfaced with a heated electrospray ionization (HESI-II) source and Orbitrap mass analyzer operated at 35,000 mass resolution. The sample extract was dried then reconstituted in solvents compatible to each of the four methods. Each reconstitution solvent contained a series of standards at fixed concentrations to ensure injection and chromatographic consistency. One aliquot was analyzed using acidic positive ion conditions, chromatographically optimized for more hydrophilic compounds. In this method, the extract was gradient eluted from a C18 column (Waters UPLC BEH C18-2.1x100 mm, 1.7 µm) using water and methanol, containing 0.05% perfluoropentanoic acid (PFPA) and 0.1% formic acid (FA). Another aliquot was also analyzed using acidic positive ion conditions; however it was chromatographically optimized for more hydrophobic compounds. In this method, the extract was gradient eluted from the same afore mentioned C18 column using methanol, acetonitrile, water, 0.05% PFPA and 0.01% FA and was operated at an overall higher organic content. Another aliquot was analyzed using basic negative ion optimized conditions using a separate dedicated C18 column. The basic extracts were gradient eluted from the column using methanol and water, however with 6.5 mM Ammonium Bicarbonate at pH 8. The fourth aliquot was analyzed via negative ionization following elution from a HILIC column (Waters UPLC BEH Amide 2.1x150 mm, 1.7 µm) using a gradient consisting of water and acetonitrile with 10mM Ammonium Formate, pH 10.8. The MS analysis alternated between MS and data-dependent MSn scans using dynamic exclusion. The scan range varied slightly between methods but covered 70-1000 m/z. Raw data files were archived and extracted as described below.

#### *WGCNA*

Weighted gene correlation network analysis (WGCNA)[4] was used to derive clusters of proteins at age 1 year based on pairwise correlations between protein features using the WGCNA package in R v4.3.0.[5] Highly correlated clusters were merged using a cut height (i.e., the Euclidean distance between clusters) of 0.3 and a soft power threshold of 6 based on iterative process to identify an optimal number of clusters. WGCNA computes a cluster membership value and associated P-value for each protein within a cluster; this value represents the degree of correlation of each individual protein feature with all other protein features within the same cluster. Following the clustering of proteins with WGCNA, protein groups (i.e., all protein features within a respective cluster) were input into the STRING database version 12.0[6] to identify common biological functions and provide names for each cluster. Protein clusters at age 1 year were applied to age 6 year protein profiling data to generate unique cluster membership and eigenvalues while maintaining identical protein groupings. Clusters were then summarized as an eigenvector based on the first principal component for each individual. This eigenvector value was utilized in subsequent statistical models to estimate associations between clusters and clinical outcomes, ‘omics, and social/environmental characteristics.

#### *STRINGdb*

The STRING database (https://string-db.org/)[6] was used to create visual representations of protein-protein interactions for each of the protein modules derived using WGCNA. For each module, individual protein features were input using the “Multiple proteins” search tool, with specification of Homo sapiens for organism. As the nature of this study was exploratory, all active interaction sources available in STRINGdb were included in network construction and visualization, including textmining, experiments, databases, co-expression, neighborhood, gene fusion, and co-occurrence; detailed explanations of these activation sources are available in the FAQ section of the STRINGdb website. Red lines indicate the presence of fusion evidence, green indicates neighborhood evidence, blue indicates cooccurrence evidence, purple indicates experimental evidence, yellow indicates textmining evidence, light blue indicates database evidence, and black indicates coexpression evidence. The default medium confidence level of 0.4 was required as a threshold for interactions; briefly, confidence levels are derived from combined scores from all sources and are representative of the level of confidence in the connection between two respective nodes.[6] All edges in the STRING network are listed in **Supplementary File S6**. Only the color of edges was meaningful for these depictions; edge lengths do not correlate to relatedness and were optimized for label clarity. Node colors do not signify meaning; colors were randomly assigned by the STRINGdb web interface.

### SUPPLEMENTARY FIGURES

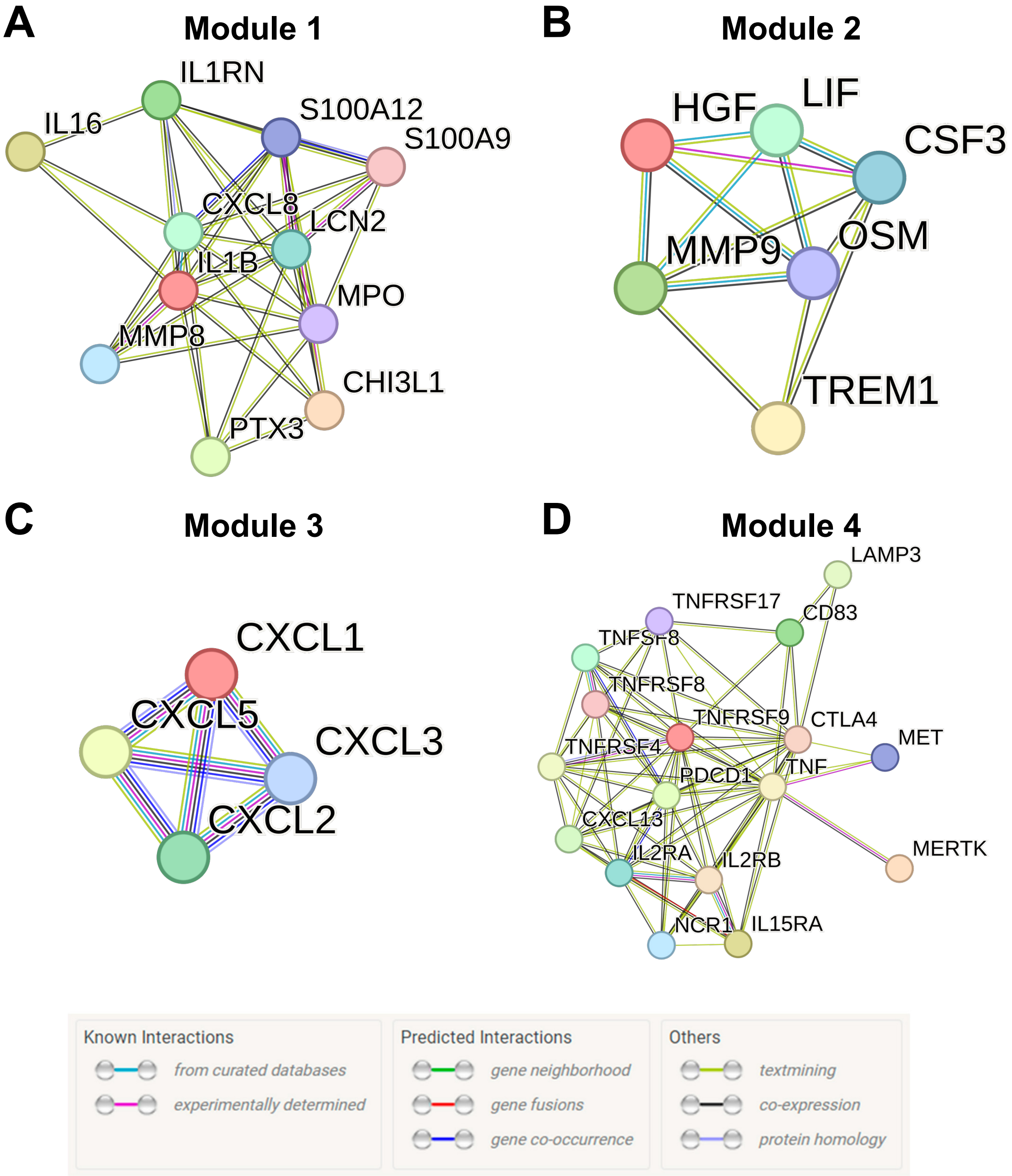


**Supplementary Figure 1. Module groupings in STRING database.** Following WGCNA, the STRING database (https://string-db.org/) was used to provide names for modules and investigate potential relationships based on known associations from literature; STRING database networks are shown for Module 1 (A), Module 2 (B), Module 3 (C), and Module 4 (D). As the nature of this study was exploratory, all active interaction sources available in STRINGdb were included in network construction and visualization, including textmining, experiments, databases, co-expression, neighborhood, gene fusion, and co-occurrence; detailed explanations of these activation sources are available in the FAQ section of the STRINGdb website. Red lines indicate the presence of fusion evidence, green indicates neighborhood evidence, blue indicates cooccurrence evidence, purple indicates experimental evidence, yellow indicates textmining evidence, light blue indicates database evidence, and black indicates coexpression evidence. Node colors are arbitrary. Edge lengths between nodes are not representative of strength of connection.

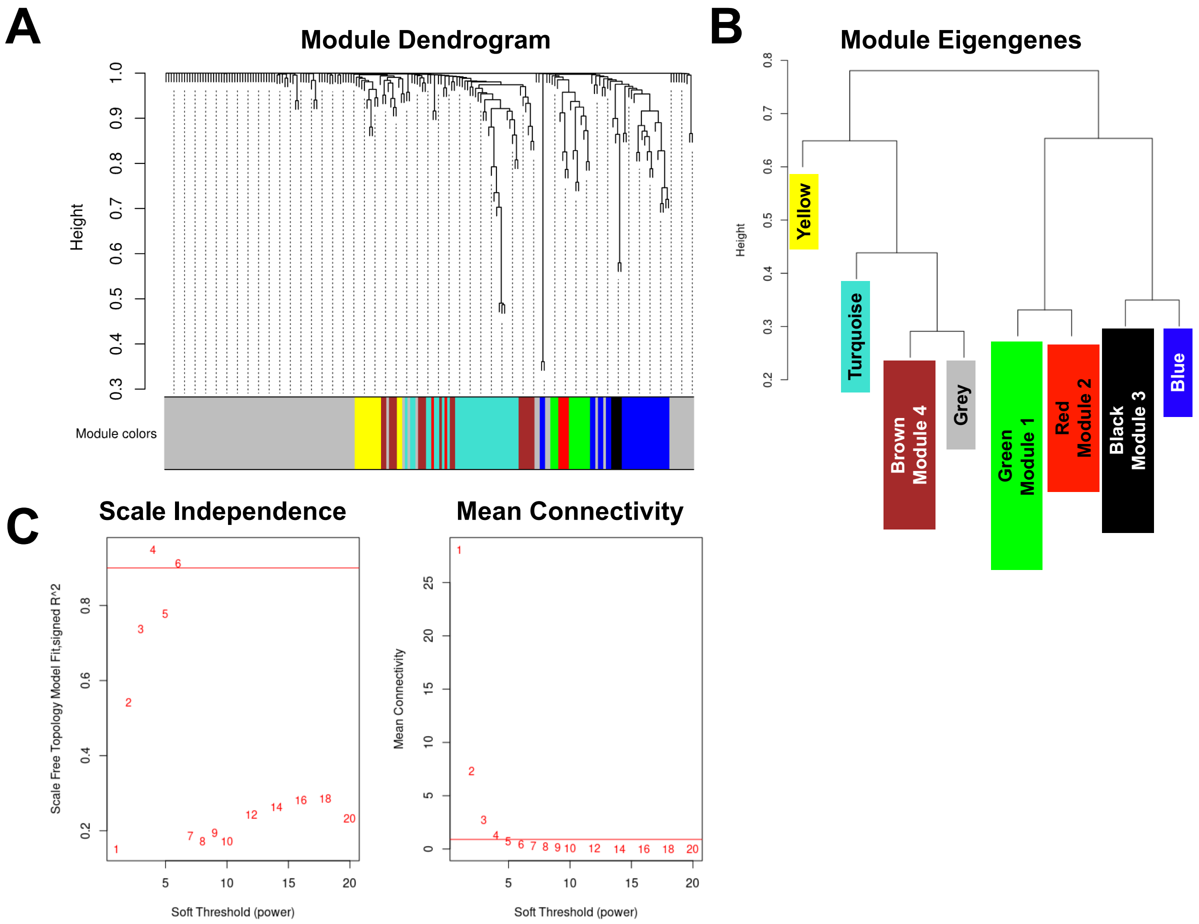


**Supplementary Figure 2. WGCNA Module Metrics.** Internal network validation metrics for the WGCNA module selection process are demonstrated, including the module dendrogram (A) demonstrating the division of proteins into one of eight modules; four of which correlated with clinical endpoints. Module eigenvalues (i.e., eigengenes) dictate the grouping of individual protein features into protein modules, shown in (B). (C-D) Scale Independence and Mean Connectivity metrics are demonstrated against soft threshold power. These metrics aid in identifying the optimal number of protein modules.

### SUPPLEMENTARY TABLES

**Supplementary Table 1. Correlation between individual protein levels at age 1 and modules.** Protein correlations determine module membership in WGCNA. Correlations between individual proteins and module eigenvalue (i.e., the first principal component) and correlation P-value are shown in the table. A total of 200 proteins were included in module analysis. A blank value under the “Module Membership” column denotes that protein was not sorted into any of the four clinically-relevant modules derived at age 1 year.

**Supplementary Table 2. Regression results for associations between module eigenvalues and clinical outcomes.** The results for regression models estimating associations between module eigenvalues at age 1 year and outcomes are demonstrated for each of the 4 modules. Beta estimates and 95% confidence intervals, P-values, and adjusted P-values to correct for multiple testing are available in base regression and fully adjusted regression models. Fully adjusted models incorporated sex, race, daycare attendance from ages 0-1 year, and duration of breastfeeding.

**Supplementary Table 3. Correlation between individual plasma protein concentrations at age 6 years and modules.** Correlations between individual proteins and module eigenvalues at age 6 years are shown in the table. A total of 200 proteins were included in module analysis. A blank value under the "Module Membership" column denotes that protein was not sorted into any of the four modules.

**Supplementary Table 4. Regression results for associations between protein modules and prenatal, perinatal, postnatal, and demographic variables.** The results for regression models estimating associations between module eigenvalues at age 1 year and social and environmental variables over the prenatal, perinatal, postnatal periods and demographic characteristics are shown. Beta estimates and 95% confidence intervals, P-values, and Adjusted P-values to correct for multiple testing are displayed in the table.

**Supplementary Table 5. Regression results for associations between individual metabolites and protein modules.** The results for regression models estimating associations between module eigenvalues at age 1 year and individual metabolites are shown. Beta estimates and 95% confidence intervals, P-values, and Adjusted P-values to correct for multiple testing are displayed in the table. Associations meeting and adjusted P-value<0.05 threshold are bolded. Regression results for individual metabolites were used as input for *MetaboAnalyst* enrichment.

**Supplementary Table 6. STRINGdb protein network edges.** The number of edges in each of the STRINGdb generated networks for module visualization are shown in this table. Protein names, identifiers, the number of edges, and module designations are shown. Module hubs based on the maximum number of edges are bolded within each module.
